# Modeling breast cancer by grafting patient tumor samples in the avian embryo: an *in vivo* platform for therapy evaluation coupled to large scale molecular analyses

**DOI:** 10.1101/2021.04.26.441511

**Authors:** Loraine Jarrosson, Clélia Costechareyre, Fanny Gallix, Séverine Ciré, Fabien Gay, Olivier Imbaud, Elisabetta Marangoni, Karine Aguéra, Céline Delloye-Bourgeois, Valérie Castellani

## Abstract

Lack of preclinical patient-derived xenograft (PDX) cancer models in which to conduct large scale molecular studies seriously impairs the development of effective personalized therapies. We report here on an *in vivo* concept consisting of implanting human tumor cells in targeted tissues of an avian embryo, delivering therapeutics, evaluating their efficacy by measuring tumors using light sheet confocal microscopy, and conducting large scale RNAseq analysis to characterize therapeutic-induced changes in gene expression. The model was established to recapitulate triple negative breast cancer (TNBC) and validated using TNBC standards of care (SOCs) and an investigational therapeutic agent.

## Introduction

The creation of innovative animal models recapitulating patient tumors as closely as possible is crucial for better understanding various types of cancer and for the development of novel therapies. Existing models struggle to establish tumors from patient biopsies in living organisms that also allow for large scale molecular analysis. Patient-derived xenografts (PDX) in the mouse have represented a breakthrough in cancer models that considerably advanced our knowledge (Dobrolecki et al., 2016). Nevertheless, they suffer from a high degree of tumor intake variability, long and variable timing of tumor growth and engraftment modality classically under the skin, thus in a context not recapitulating the tumor microenvironment (Shi et al., 2020). All of these challenges prevent the use of such models in extensive large-scale studies comparing functional responses of single patient tumors to multiple treatment regimens and correlation with specific molecular signatures.

Additionally, such limitations of existing preclinical models curtail the investigation of molecular and therapy-response heterogeneity among patients suffering from the same types of cancers, which investigations would otherwise usher in the development of personalized medicine. Patients would thus benefit substantially from paradigms that afford rapid prediction of their responsiveness or resistance to proposed therapies. This is of central interest for those suffering from cancers with poor outcomes, for which the choice of treatment regimens is particularly complex and fraught with risk.

Breast cancer is the second-leading cause of death in women worldwide. The triple negative breast cancer (TNBC) subtype represents 10–15% of all diagnosed breast cancers, and has the poorest prognosis with a median overall survival for metastasized patients of approximately eighteen months (Hwang et al., 2019; Mehanna et al., 2019). TNBC usually appears as a high-grade ductal carcinoma, defined by the lack of expression of estrogen receptor (ER), progesterone receptor (PR), and HER2 receptor, hence the term “triple negative”, and often accompanied by distant metastases. Extensive molecular profiling refined the classification of the different breast cancer subtypes, revealing heterogeneity reflected by six different molecular profiles (Lehmann et al., 2011). Tumor sequencing also supported efforts to produce personalized or patient-tumor-specific treatments, but finding a clear correlation between sequence profiles and therapeutic efficacy remains elusive. Patients with TNBC typically receive chemotherapy; however, recurrence rate remains high and the development of chemotherapy resistance occurs frequently. Further still, the recurrence rate remains high and the development of chemotherapy resistance occurs frequently. And finally, dissemination of TNBC is a major clinical issue, particularly considering that the predominant lung, brain, and bone metastases remain incurable (Medina et al., 2020; Vagia et al., 2020). Accordingly, the development of novel therapeutic options for TNBC continues to be a high priority.

A variety of murine models, based either on engraftment of breast cancer cells or on genetically-engineered tumorigenesis, has been developed to elucidate the mechanisms underlying tumor development, response to therapy and acquired chemotherapy resistance (Park et al., 2018). Despite the utility of these models, comprehensive studies modeling patient tumor heterogeneity for biomarker discovery are lacking. To address this unmet need, grafting of tumor cells onto the chorioallantoic membrane of the avian egg has been considered as an alternative of murine xenografts. However, despite the advantages of such a simple *in vivo* model, tumor growth varies substantially in the extra-embryonic environment, in stark contrast to the growth observed in the context of a tumor-infiltrated organ.

Seeking to overcome the foregoing limitations, we recently reported on the successful modeling of pediatric neuroblastoma in the avian embryo (Delloye-Bourgeois et al., 2017). To more accurately mimic the *in vivo* milieu, we grafted neuroblastoma cells within tissues expected to provide a microenvironment representative of their cellular origin. This strategy proved highly effective in recapitulating key aspects of the disease, revealing that the embryonic environment provides a relevant context in which to model tumor growth and dissemination. Indeed, mechanisms at work during development and tumorigenesis have emerged over years as being closely related. For example, numerous studies have reported physiopathological conditions in which cancer cells take advantage of signaling pathways such as Wnt, Sonic hedgehog (SHH), TGF or BMP, acknowledged for their key contributions to embryonic development (Wakefield and Hill, 2013).

In breast cancer, this phenomenon manifests itself in the form of molecular communication between tumor cells and bone stroma that lie at the core of metastasis progression (Sethi and Kang, 2011). The bone microenvironment consists of mineralized extracellular matrix and various resident cell types, including osteoblasts, osteoclasts, mesenchymal stem cells, bone marrow endothelial cells, hematopoietic cells, and adipocytes. Bone homeostasis relies on the coordinated activity of osteoclasts and osteoblasts resorbing and renewing bone matrix, respectively. Such renewal processes are analogous to bone formation in the developing embryo, and perhaps unsurprisingly, involve developmental signaling pathways as attested by multiple studies (Mukherjee et al., 2018; Sethi and Kang, 2011; Wakefield and Hill, 2013; Ye and Jiang, 2016). For example, CXCR4/CXCL12 signaling plays a prominent role in several processes during embryogenesis, as demonstrated by the phenotypes resulting from ligand and receptor gene deletions (Kawaguchi et al., 2019; Nagasawa et al., 1996), and also controls key processes of bone physiology in adults (Kawaguchi et al., 2019). Interestingly, cancer cells establishing metastases in bone participate in these molecular exchanges, exploiting secreted signals normally reserved for osteoblasts and hematopoietic stem cells (Hiraga, 2019). Likewise, it has been recently established that upregulated CXCR4 expression enables some cancer cells to metastasize to bone (Coniglio, 2018; Hiraga, 2019). Once there, CXCR4 exerts a pro-osteolytic effect.

These parallels between embryonic development, homeostasis, and tumorigenesis led us to hypothesize that engrafting patient-derived tumor cells in embryonic regions committed to form tumor-relevant tissues would yield a microenvironment supportive of tumor formation. In this study, we report the creation of an *in vivo* model encompassing the generation and analysis of miniaturized replicas of patient tumors in targeted regions of the chicken embryo and their evaluation in preclinical studies. Breast cancer cells were implanted into a selected embryonic region, which was committed to form bones, to mimic a prominent metastatic site of breast cancer. We show using both the MDA-MB-436 TNBC cell line and patient biopsies that our avian model is remarkably effective in promoting rapid tumor intake (e.g. about 24 hours), even when the initial number of cells is small. We also report the establishment of intravenous administration of standard therapies and the analysis of their efficacy on a series of patient tumor replicas. We further demonstrate that our patient tumor replicas allow for the coordinated evaluation of therapy-induced changes in both tumor genotype and phenotype. Finally, the exploitation of our innovative model for preclinical investigations was validated using L-Asparaginase (ASNase), an active substance of a therapy currently being evaluated in several clinical trials (e.g. Erytech’s Trybeca-1 and Trybeca-2, evaluating the efficacy of red cell-encapsulated ASNase against pancreatic cancer and TNBC respectively). Altogether, our study establishes the patient-derived xenograft avian model (AVI-PDX™) as an innovative tumor model for preclinical programs, which opens new avenues for the evaluation of mono and combination therapies and the characterization of signatures predictive of patient responses.

## Results

### Micrografting breast cancer cells in the developing somites of the avian embryo

To design a model of tumorigenesis mimicking bone metastasis, we thought to target particular somites of the developing avian embryo. Somites are transient bilateral epithelial structures, each having a spheroidal shape. They are generated within the paraxial mesoderm (also referred as the presomitic mesoderm) according to a metameric pattern along the rostro-caudal body axis of the embryo (Maschner et al., 2016). Somites provide the basis for the vertebral column and ribs, also giving rise to bone and trunk musculature derivatives (Williams *et al*, 2019). Over the years, the chick embryo has become a reference model and its extensive use has contributed significantly to major advances in our understanding of developmental biology in vertebrates. Accordingly, somitogenesis has been very well documented, including extensive reporting on spatial and temporal hallmarks, which allowed us to appropriately place patient-derived tumor cells during the grafting procedure (Berti et al., 2015; Pourquié, 2018).

Furthermore, chicks and humans have similar numbers of somites, while mice have about 10 additional pairs. Newly generated somites, staged I-II according to Christ and Ordahl, are epithelial spheres with a central cavity, the somitocoel, filled by mesenchymal cells. From stage III, the ventral mesenchymal compartment becomes distinct from the dermomyotome dorsal one. Ventral cells dissociate through epithelio-mesenchymal transition to form the sclerotome, and begin to express specific regulator genes such as *pax1*, and thereafter *pax9*, under the influence of morphogens released by surrounding floor plate and notochord tissue organizers, as well as by the surface ectoderm and the neural tube. In the chicken embryo, the first somite pair is visible at stage 7 according to Hamburger and Hamilton staging (HH7, 23-26 hours post-gestation) (Hamburger and Hamilton, 1992). During stage HH14 (50-53 hours post-gestation), a stage that we had previously found convenient for grafting procedures (Delloye-Bourgeois et al., 2017), 22 somite pairs are formed. Also, around this time, somite compartmentalization has already occurred for the oldest somites, including ventral cells progressing towards sclerotome differentiation under the control of various signals.

We used a well-characterized TNBC cell line, MDA-MB-436, to determine whether this presumptive skeletal embryonic region could support survival and growth of breast cancer cells. Cells were labeled with the vital fluorochrome CFSE, and systematically grafted within somites 12 and 13 of multiple HH14 embryos. Forty-eight (48) hours later, HH25 embryos were collected (**Figure 1A**). Systematic fluorescence detected with a stereomicroscope was indicative of the presence of MDA-MB-436 cells in 100% of grafted embryos (**Figure 1B**). We took advantage of confocal light sheet microscopy to establish a procedure for analysis of tumor cells at the whole organism level. Embryos were fixed and subsequently cleared using the Ethyl Cinnamate (ECi) procedure. Notably, we consistently observed the formation of tumor masses within the developing somites, with a few cells escaping from these dense masses (**Figure 1B**). We then set up an analytic pipeline to precisely measure the volume occupied by tumor cells (**Figure 1B**). Tumor sizes suggested that MDA-MB-436 cells proliferated within the somitic environment. We further performed a Ki67 immunolabeling on cryosections of the embryos, which confirmed a high mitotic index of grafted MDA-MB-436 cells, with a mean of 34% of Ki67^+^ cells among analyzed cryosections (n=10) (**Figure 1C**). Thus, our model induces TNBC cells to take root within the developing somitic environment, to proliferate, and to form measurable tumor masses with ubiquitous and reproducible tumor intake occurring in no more than 48 hours.

**Figure 1:**
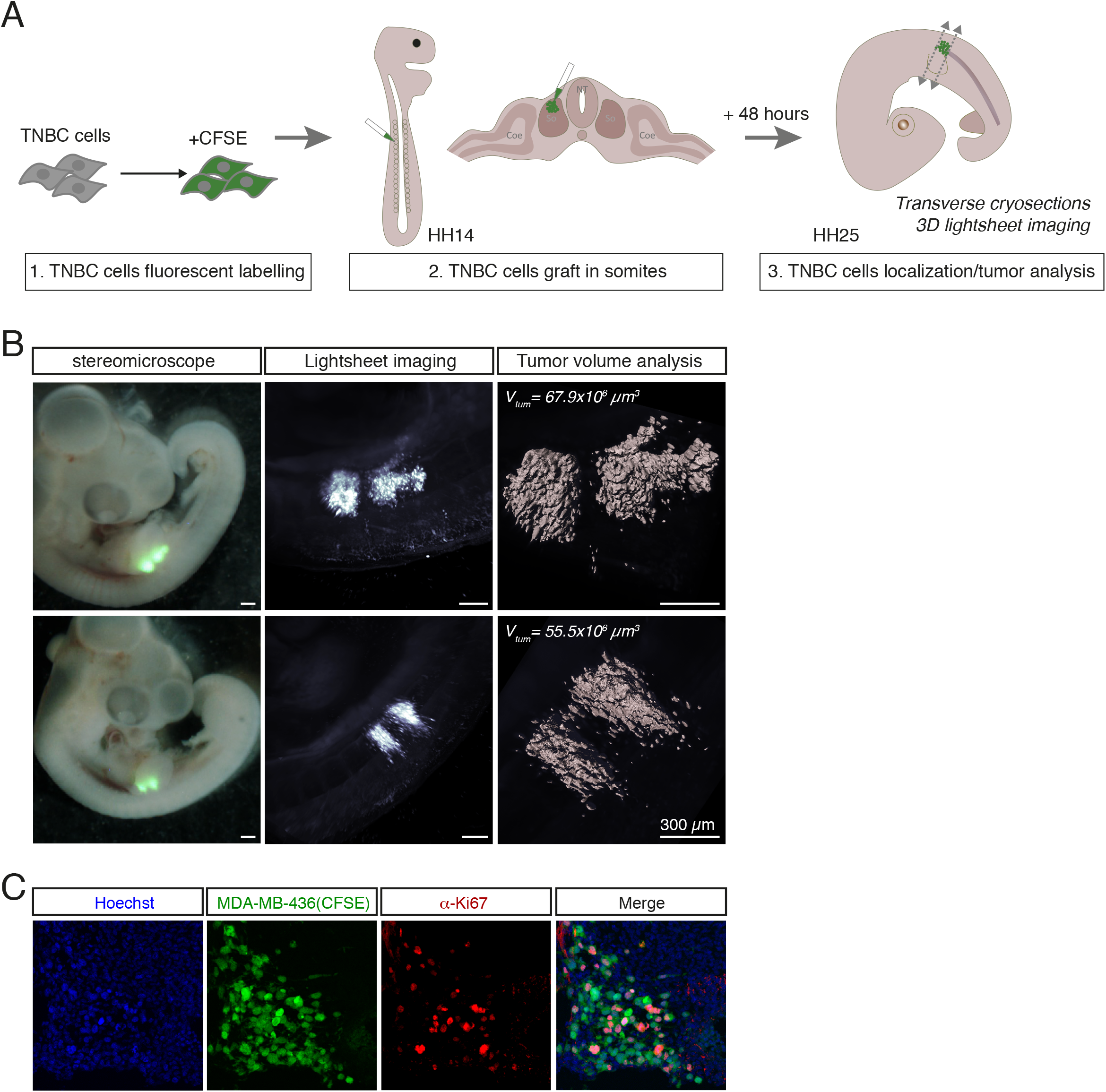
Grafting of MDA-MB-436 TNBC cells within developing somites allows tumor formation and proliferation. **A**. Set up of TNBC cell grafting in the developing somites: TNBC cells are labeled with CFSE and microinjected in the developing somites at HH14 stage. Grafted embryos are harvested for imaging analysis 48 hours after the graft, at HH25 stage. **B**. Detection and volumetric analysis of TNBC tumor masses formed 48 hours after the graft: TNBC fluorescent masses are detected with a stereomicroscope (left panels), and next imaged in whole cleared embryos using lightsheet imaging (middle panels), which allows a precise quantification of tumor volumes using Imaris 3D software (right panels). **C**. Detection of Ki67+ proliferating cells by immunofluorescence (in red) on cryosections of MDA-MB-436 tumor masses (in green, CFSE+) formed in HH25 chick embryos. Chick and human nuclei are labelled with Hoechst (in blue). Scale bars: 300 µm.

### Administration of SOCs in the avian model recapitulates TNBC response to standard anticancer therapies

We next wanted to understand whether our tumor model would respond to current anticancer therapies similarly to how patient tumors respond *in vivo*, with the goal of aiming to create a new model relevant for preclinical studies. We chose to work with two major standards of care (SOCs), gemcitabine and carboplatin, commonly used in the treatment of TNBC.

First, we set up a procedure to determine optimal doses of drugs. Having access to the vascular network of the chorioallantoic membrane that irrigates the developing embryo, we performed intravenous (IV) injections of increasing doses of gemcitabine or carboplatin, in multiple HH20 chick embryos (approximately 72H post-gestation) (**Figure 2A**). To determine the maximum tolerated dose (MTD), we established a list of criteria that we examined 24 hours post-injection (HH25 embryos) including: survival of the embryo, morphological checkpoints, and global growth by measuring the body surface area (BSA), as detailed in the methods section (**Figure 2A**). A survival rate below 75% was indicative of dose toxicity, excluding further examination of the concerned group. The correct stage-related development of embryos was systematically assessed by checking their craniofacial morphology (presence of each cerebral compartment and eyes), the presence of four limb buds, their cardiac morphology, and the anatomy of embryonic annexes such as the allantois. For gemcitabine, we observed that doses higher than 17.1 mg/kg induced massive embryonic death. Doses between 1.37 and 3.42 mg/kg were associated with a survival rate above 75%, but significantly affected global growth (as measured by the BSA criteria) and morphological checkpoints. We thus fixed the gemcitabine MTD at 0.68 mg/kg in chick embryos (**Figure 2B**). Following the same procedure, the carboplatin MTD was determined to be 92.3 mg/kg (**Figure 2C**). As gemcitabine and carboplatin are frequently used as a combi-therapy, simultaneous injection of gemcitabine and carboplatin MTDs was tested in multiple HH20 chick embryos (**Figure 2D**). We could observe that the co-treatment was perfectly tolerated *in ovo*, according to the criteria listed in this section.

**Figure 2:**
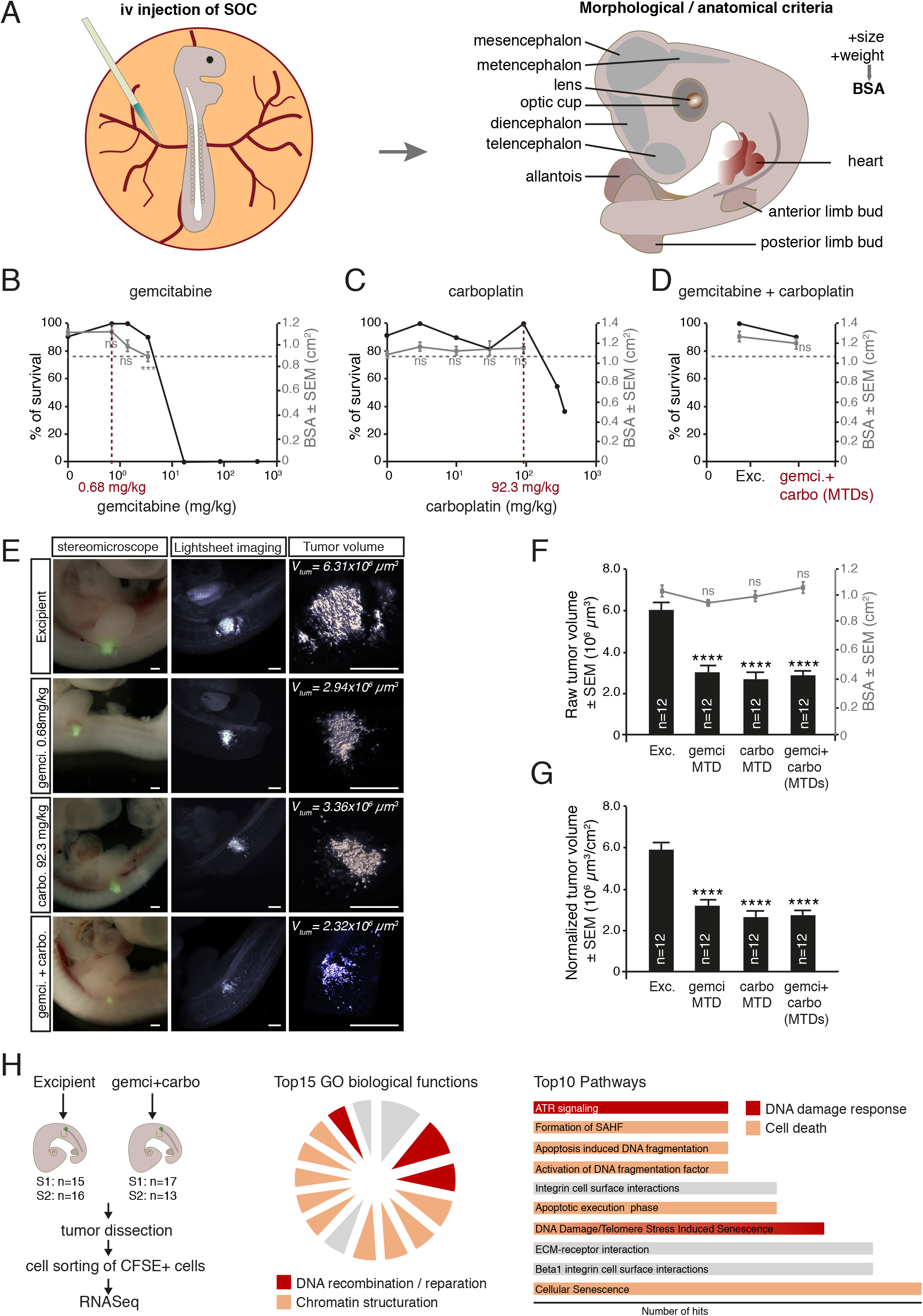
Administration of SOCs in the avian model efficiently triggers anticancer response of grafted TNBC cells. **A**. Schematic representation of the procedure of intravenous SOC administration in the avian model (left panel) and illustration of morphologic, anatomical and morphometric criteria analyzed for each embryo after drug administration (right panel). **B-D**. Survival rate (left axis) and mean body surface area (BSA, right axis) of chick embryos injected with increasing doses of gemcitabine (**B**), carboplatin (**C**) or combination of both maximum tolerated doses (MTDs) (**D**). Each dose was administered to a minimum of 10 embryos, using excipient (NaCl) as a control. The MTD of gemcitabine and carboplatin was defined as the higher dose of drug associated with a survival rate higher than 80% and a mean BSA similar (-ie, non-statistically different) from embryos treated with NaCl. MTDs are indicated in red on the abscissa axis. Error bars indicate SEM. **E-G**. Analysis of tumor growth by 3D lightsheet imaging and tumor volumetric analysis (**E**) of chick embryos treated with gemcitabine MTD (n=12), carboplatin MTD (n=12), a combination of both (n=12), or excipient (n=12) in series of chick embryos grafted with MDA-MB-436 cells. Scale bar: 300 µm. Mean raw tumor volumes and mean BSA for each experimental condition was measured (**F**). Mean tumor volumes normalized on mean BSA are also reported to take into account slight variability of embryonic growth (**G**). Error bars indicate SEM. ****: p<0.0001. ns: non-significant. **H**. Whole RNASeq analysis of MDA-MB-436 cells sorted from series of grafted chick embryos treated either with excipient or with a combination of gemcitabine and carboplatin MTD. The procedure is illustrated in the left panel and the number of embryos for each of the two replicates is indicated. Ninety-one (91) differentially expressed transcripts (padj < 0.05 and fold change >1.5) and 111 “ON/OFF” transcripts were considered for global analysis of biological functions (middle panel) and signaling pathways (right panel) concerned by gene expression change. Within the 15 most significantly regulated biological functions, the ones related to DNA recombination/reparation and chromatin structuration are highlighted on the diagram. Pathways related to DNA damage response and/or cell death are highlighted within the top 10 significantly regulated pathways.

We next assessed whether administration of gemcitabine and carboplatin MTDs, alone or in combination, could impact on MDA-MB-436 tumor growth in the avian embryo. TNBC cell line was described to be sensitive to both SOCs in a range of *in vitro* and *in vivo* studies (Larsson et al., 2020; Mintz et al., 2020). Each SOC MTD was injected intravenously in randomized batches of avian embryos 24 hours after engraftment of MDA-MB-436 cells. 24 hours after SOC administration, embryos were harvested and tumor volumes were analyzed by light sheet microscopy and subsequent 3D image analysis, as described in Figure 1 (**Figure 2E**). Administration of gemcitabine and carboplatin alone or in combination induced a significant decrease in mean tumor volume, without affecting the global embryonic growth (**Figures 2F and 2G**). While each SOC alone drastically affected tumor growth, simultaneous administration of both did not trigger any additive effect on MDA-MB-436 tumors as compared to single SOC administrations. Thus, within a brief window of time, 24 hours, our avian model of breast cancer cell grafting within the developing somites of chick embryos recapitulates tumor growth and tumor response to SOCs.

### RNAseq analysis of gemcitabine/carboplatin-triggered transcriptional regulations in tumors formed in the avian embryo

Next, we asked whether we could combine our *in vivo* tumor model with large scale analyses to evaluate the impact of SOCs on TNBC cells at the molecular level. We micro-dissected the tumors from a series of grafted embryos and treated them with either the excipient or with a combination of gemcitabine and carboplatin. For each condition, tumor cells embedded in the chick embryonic tissues were dissociated and sorted for bulk RNA Sequencing (RNAseq) (**Figure 2H**). Given the small number of replicates (duplicates for each condition), we chose to select stringent parameters to perform the differential expression analysis, to avoid false positive findings, as described in the methods section. Using these criteria, the analysis revealed a set of 91 transcripts significantly regulated in chemo-treated tumors as compared to excipient (padj<0.05, fold-change>1.5) and 111 transcripts whose expression was detected only in one of the two experimental conditions (“ON/OFF” transcripts, as explained in the methods section). This list of 202 transcripts was further investigated with toppgene software (https://toppgene.cchmc.org) to assign key biological functions and signaling pathways impacted by gemcitabine/carboplatin treatment. Notably, among the 15 top biological functions represented, 12 were related to DNA repair / recombination processes and to the regulation of chromatin structure (**Figure 2H**). Similarly, among the top 10 signaling pathways associated with gene expression regulation upon gemcitabine/carboplatin treatment, 7 were directly related to DNA-damage response, apoptosis and senescence induction (**Figure 2H**). Interestingly, these findings are in accordance with well-described modes of action of gemcitabine and carboplatin in the clinic, which include the targeting of DNA replication and repair mechanisms, leading to activation of the DNA-Damage Response and subsequent death of chemo-sensitive cells.

### Adaptation of the micrografting technique to breast cancer patient samples: creation of an avian Patient-Derived Xenograft (AVI-PDX™) model

To further extend our model, we next thought to test whether such a micrografting technique could be adapted to fresh or frozen patient samples, without any intermediate culture step potentially altering the tumoral features. We selected 15 different patient samples with diverse origins and preparation modalities: “rough” patient samples without any experimental handling (n=11), patient samples derived from murine PDX models (n=4); fresh (n=6) and DMSO-cryopreserved (n=9) samples; samples originating from primary tumors (n=10) or brain metastases (n=5) (**Figure 3A**). Each sample was dissociated and the global cell content was labeled with CFSE prior to engraftment. The tumor establishment rate 48 hours after the implantation was remarkably above 70% for all samples tested except for OF-BRE-007 for which it was of 34%. While tumor size 48 hours after engraftment was homogeneous among embryos engrafted with the same sample cell content, mean tumor volume for each sample varied between 7.13×10^3^ and 2830×10^3^ µm^3^, indicating heterogeneity in tumor sample cell content and/or behavior within avian embryonic tissues (**Figures 3A and 3B**). Thus, we were able to generate 14 avian PDX (AVI-PDX™) models from breast cancer patient samples, with homogeneous tumor growth among embryonic replicates and with an exceptionally fast and high rate of tumor intake. We tested the effect of gemcitabine in combination with carboplatin in 4 AVI-PDX™ models obtained from 4 rough TNBC patient samples, 2 primary tumors and 2 brain metastases (**Figure 3C**). Comparison of tumor volumes between excipient and SOC-treated embryos revealed that two AVI-PDX™ models (OF-BRE-002 and OF-BRE-nbt311) did show a significant anticancer response to SOC, while the two others (OF-BRE-012 and OF-BRE-nbt783-derived AVI-PDX™) were completely resistant to SOC administration. Thus, tumor replicas from different patients exhibited response heterogeneity, which is a hallmark of actual TNBC patient tumors in the clinic.

**Figure 3:**
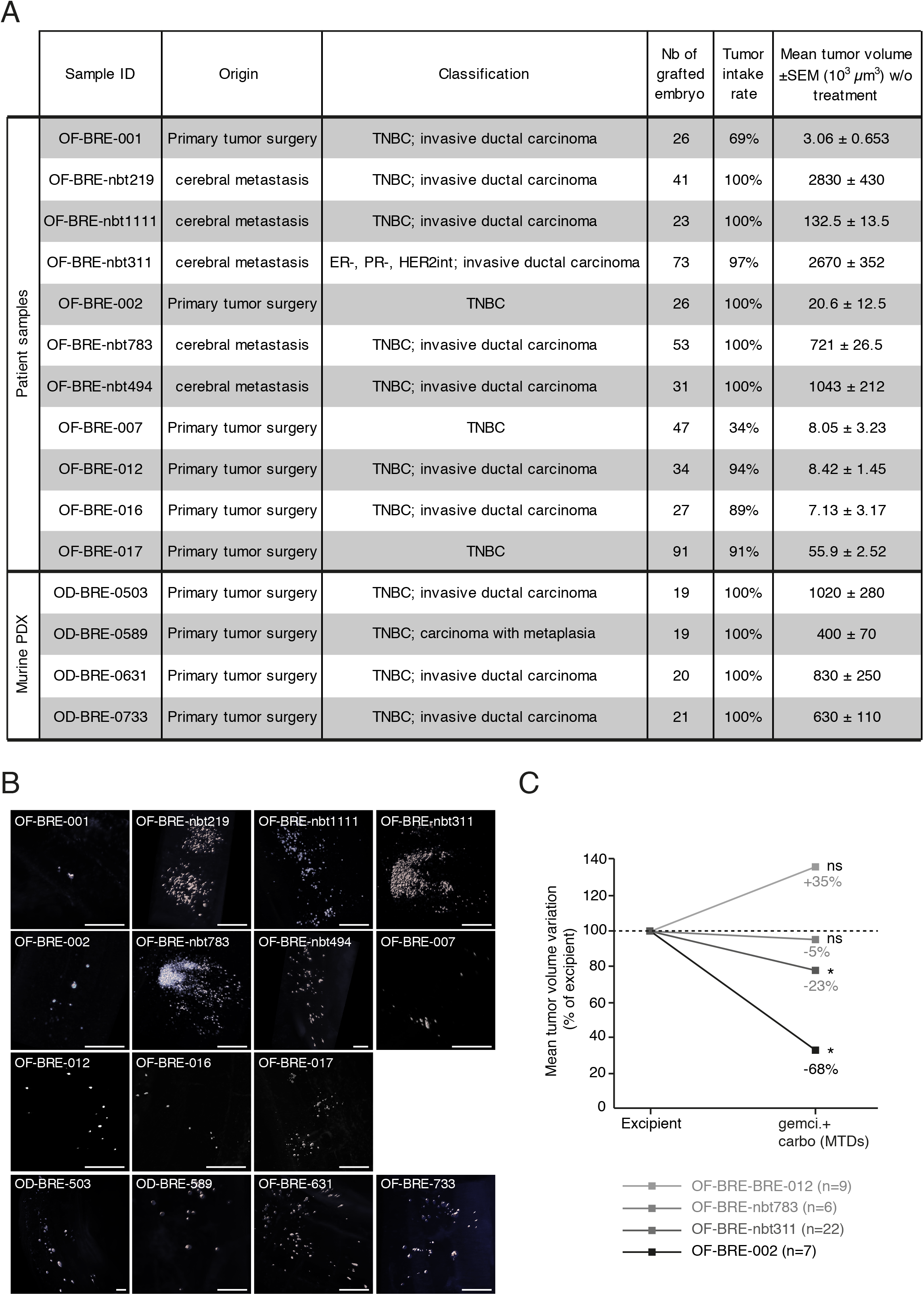
The avian model is adapted to the micrograft of patient samples. **A**. Table recapitulating breast cancer sample characteristics – origin, classification – and their behavior in the avian approach – tumor intake rate, mean tumor volume as measured by 3D lightsheet imaging and subsequent volume measurement-. **B**. Representative 3D lightsheet images obtained with each patient sample referenced in the A. Scale bar: 300 µm. **C**. Effect of gemcitabine/carboplatin administration in series of embryos engrafted with 4 different patient samples, as compared to excipient administration. Tumor volumes were measured using lightsheet microscopy and 3D image analysis; results are presented as mean tumor volume variation in gemcitabine/carboplatin-treated embryos as compared to excipient treated embryos. The number of embryos analyzed for each patient sample is indicated on the graph.

### Preclinical evaluation of L-Asparaginase efficacy in TNBC treatment using the AVI-PDX™ system

These results indicated that the AVI-PDX™ model might be highly relevant and valuable for evaluating the efficacy of candidate therapeutic compounds. L-Asparaginase (ASNase) hydrolyzes L-asparagine (ASN) and Glutamine (GLN) into aspartic (ASP) and glutamic (GLU) acids and ammonia, leading to ASN and GLN removal from the circulation, which in turn causes metabolic dysfunction, cell cycle arrest, apoptosis and tumor starvation. ASNase is routinely used in the treatment of some hematological malignancies, most notably acute lymphoblastic leukemia (ALL) (Ahlke et al., 1997; Estlin et al., 2000; Müller and Boos, 1998).

To assess the impact of ASNase on tumor cell survival *in vitro*, we first conducted experiments with the MDA-MB-436 TNBC cell line (**Figure 4A**). The objective was to evaluate the *in vitro* sensitivity of this cell line to recombinant *E*.*coli* ASNase alone by determination of the concentration of drug that gives a 50% inhibition of cell viability (IC50). Although cytotoxicity curve profiles showed some heterogeneity, the IC50 determination remained reproducible between experiments. On Day 4, a dose-response and a similar plateau effect were observed for doses of ASNase from 1 to 10 U/mL. However, 10% of cells still remained viable at even the highest concentration tested (10 U/ml). The mean IC50 of ASNase from the 3 experiments on MDA-MB-436 cells after a 4-day exposure was determined to be 0.19±0.06 U/mL.

**Figure 4:**
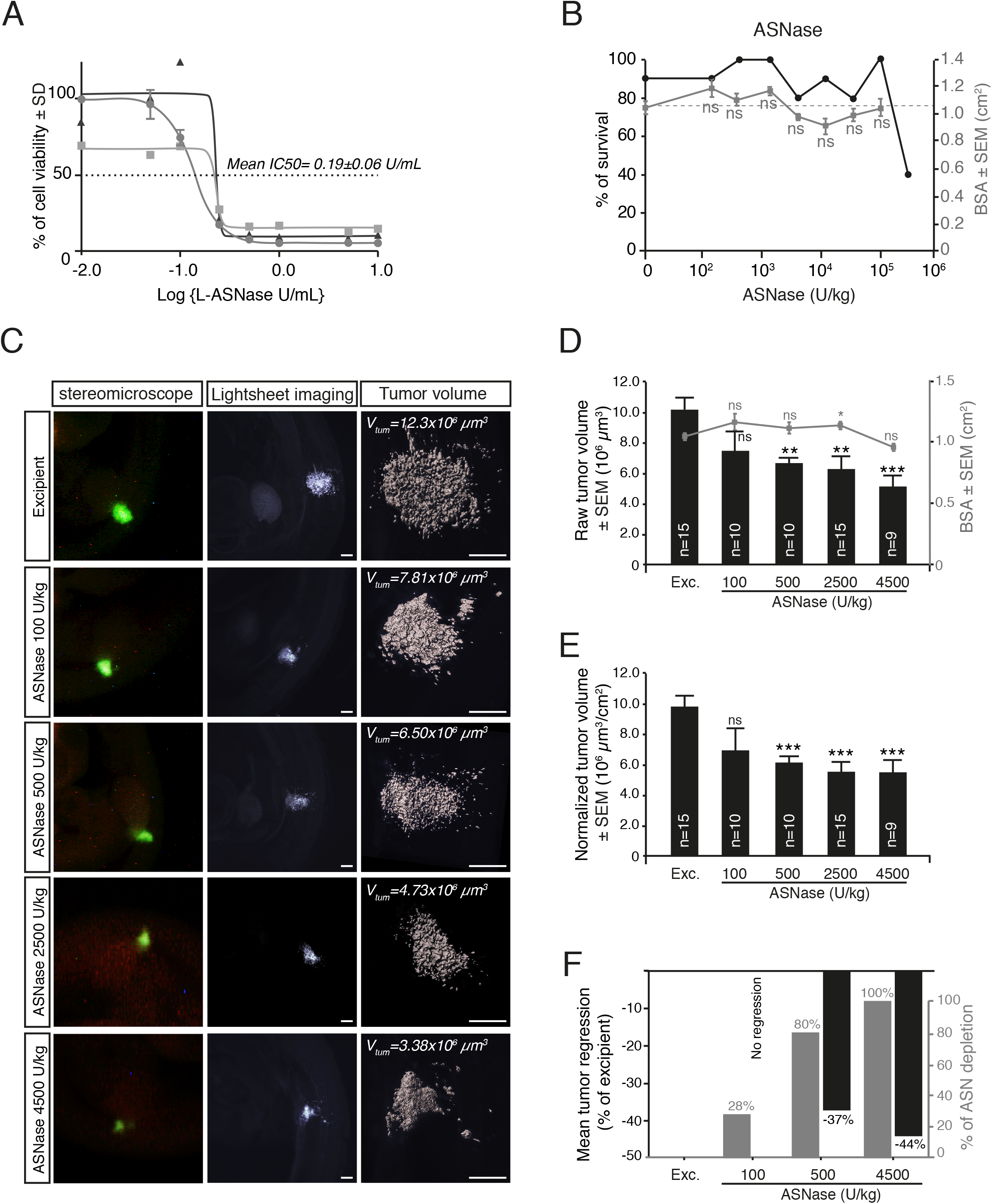
Preclinical evaluation of L-Asparaginase efficacy in TNBC using the avian model. **A**. *In vitro* evaluation of MDA-MB-436 cell viability, after treatment with increasing doses of ASNase. 3 experimental replicates are presented, allowing to estimate a mean IC50 of 0.19±0.06 U/mL. **B**. Survival rate (left axis) and mean body surface area (BSA, right axis) of chick embryos injected with increasing doses of ASNase. Each dose was administered to a minimum of 10 embryos, using excipient (NaCl) as a control. Error bars indicate SEM. **C-E**. Analysis of tumor growth by 3D lightsheet imaging and tumor volumetric analysis (**C**) of chick embryos 24 hours after treatment with increasing doses of ASNase or excipient in series of chick embryos grafted with MDA-MB-436 cells. The number of embryos analyzed for each experimental condition is indicated on the graph. Scale bar: 300 µm. Mean raw tumor volumes and mean BSA for each experimental condition was measured (**D**). Mean tumor volumes normalized on mean BSA are also reported to take into account slight variability of embryonic growth (**E**). Error bars indicate SEM. **: p<0.01; ***: p<0.001. ns: non-significant. **F**. Quantification of plasma ASN in series of chick embryos 24 hours after iv injection of increasing doses of ASNase as compared to excipient, to estimate ASN reduction rate. A minimum of 10 embryos per condition were used to harvest blood samples. Reduction rates are presented together with the corresponding tumor regression rates obtained in the avian model grafted with MDA-MB-436 cells.

Next, we measured ASNase tolerance in the avian embryo to estimate its maximum tolerated dose (**Figure 4B**). Intravenous injection of increasing doses of ASNase (150 U/kg to 1.22×10^5^ U/kg) in E3 chick embryos affected neither embryonic survival, morphogenesis, nor global growth rate, reported by the body size area (BSA). At doses of 3.57×10^5^ U/kg and higher, embryo survival was compromised and largely fell below the 75% cutoff, indicating toxicity. Interestingly, 4500 U/kg corresponds to a high dose in the avian embryo model and is quite below the total dose currently given in the clinic in repeated administration (superior to 10 000U/kg). We thus chose to fix 4500 U/kg as the maximum dose for further preclinical evaluation of ASNase efficacy.

We then evaluated the antitumor efficacy of increasing doses of ASNase, ranging from 100 to 4500 U/kg, in series of chick embryos engrafted with MDA-MB-436 cells. As set up for SOCs, a single administration of ASNase was performed 24 hours after the graft, and treated embryos were harvested 24 hours later for tumor growth analysis (**Figures 4C-4E**). While the lowest dose (100 U/kg) did not affect tumor volume (neither raw nor embryonic growth rate-normalized volume), higher doses each triggered a significant decrease in mean tumor volume without affecting global embryonic growth. Notably, in the MDA-MB-436 avian model, a slight dose effect of ASNase was observed with a mean decrease in raw tumor volume, increasing from 34% at 500 U/kg to 49% at 4500 U/kg, and from 37% to 44% when the tumor volumes were normalized to embryonic growth. As ASNase acts by depleting cells of available ASN, we thought to examine whether these efficacy experiments correlated with plasma ASN levels in treated embryos. Blood was harvested from a series of embryos that received the different doses, for evaluation of plasma amino-acid levels by HPLC (**Figure 4F**). We confirmed that increasing doses of ASNase administrated intravenously triggered a dose-dependent reduction of plasma ASN, ranging from 28% of excipient at 100 U/kg to complete reduction at 4500 U/kg. At 500 U/kg, ASN was depleted by 80%. These reduction rates mirrored the antitumor efficacy of ASNase, which was insignificant at 100 U/kg and increased from 500 to 4500 U/kg (Figures 4C-4E). Thus, ASNase enzymatic activity is effective and quantifiable in the avian embryo system, and its activity impedes MDA-MB-436 tumor growth in a dose-dependent fashion.

These first results encouraged us to further evaluate ASNase in AVI-PDX™ models, with the aim to set this candidate therapy against TNBC patient heterogeneity (**Figure 5A**). We worked with 500 and 4500 U/kg doses of ASNase that were previously shown to trigger both satisfactory ASN and MDA-MB-436 tumor volume reductions in the avian embryo. Fourteen (14) AVI-PDX™ models were engineered, among which 9 showed an anti-tumor response to ASNase with at least one of the two doses tested (green samples in **Figure 5A**). Four of these AVI-PDXTM models were achieved on limited number of embryos, excluding the possibility of performing statistical analysis. Thus, we focused on the remaining 10 models and our statistical evaluation confirmed that 60% of AVI-PDX™ responded significantly to ASNase treatment, irrespective of the origin of the patient sample (**Figure 5C-5F**).

**Figure 5:**
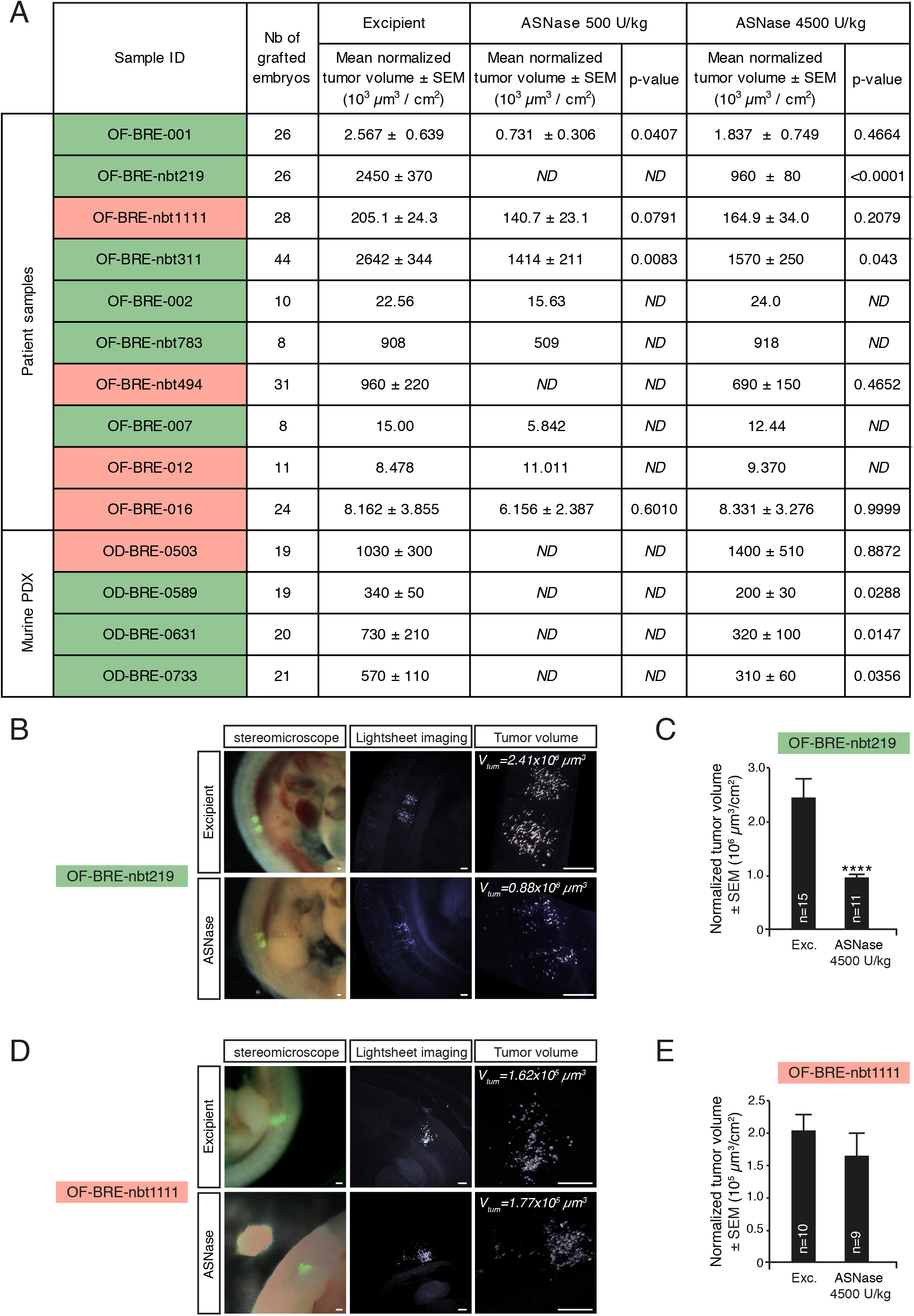
Preclinical evaluation of L-Asparaginase efficacy in the TNBC AVI-PDX model. **A**. Table recapitulating the effect of ASNase administration (500 and/or 4500 U/kg) on mean tumor volume (as compared to excipient) for each of the 14 patient samples grafted in the AVI-PDX system. Patient samples showing an anti-tumor response to ASNase are highlighted in green (9 out of 14 samples), while non-responder patient samples are highlighted in red (5 out of 14 samples). ND: Not Determined. **B-E**. Analysis of tumor growth by 3D lightsheet imaging and tumor volumetric analysis (**B**,**D**) of chick embryos 24 hours after treatment with increasing doses of ASNase or excipient in series of chick embryos grafted with OF-BRE-nbt219 (responder, **B**,**C**) or OF-BRE-nbt1111 (non-responder, **D**,**E**) TNBC samples. Scale bar: 300 µm. Mean tumor volumes normalized on mean BSA are reported for each patient samples in excipient- and ASNase-treated series of embryos (**C**,**E**). The number of embryos analyzed for each experimental condition is indicated on the graph. Error bars indicate SEM. ****: p<0.0001.

### Implementation of breast cancer micrografting technique in the avian embryo: targeting the developing brain to model cerebral metastasis

Lastly, we investigated whether we could develop an avian tumor model recapitulating another metastatic microenvironment relevant to TNBC pathology.

Metastasis to the brain is one of the hallmarks of aggressive breast cancer that appreciably affects disease outcome (Karginova et al., 2015). The brain microenvironment promotes the adaptation of highly specialized cancer cells and the formation of tumors possessing unique characteristics. Moreover, the ability of therapeutics to penetrate the blood-brain barrier continues to present a daunting challenge. Therefore, having a PDX model that could efficiently and robustly reproduce brain metastases would provide a complementary and powerful tool for the preclinical evaluation of candidate therapeutics. We engrafted either a TNBC cells (MDA-MB-436) or 1 of 7 patient samples obtained from murine PDX, directly in the brain parenchymas of a series of HH14 chick embryos (**Figure 6A**). The MDA-MB-436 cell line grafted efficiently, leading to a very high rate of tumor intake rate (93%) 48 hours after engraftment. Analysis of grafted embryos by confocal light sheet microscopy confirmed the formation of tumor masses within the brain parenchyma of all embryos (**Figures 6B and 6C**). Moreover, a Ki67 immunolabeling performed on cryosections of grafted embryos revealed a high mitotic index of grafted MDA-MB-436 cells, with a mean of 41% (± 6.4%) of Ki67^+^ cells (n=4 fields) (**Figure 6D**). These data confirmed that TNBC cells are able to form tumors within the developing chick brain tissue and to continue proliferating. Patient samples obtained from 7 murine PDX (5 Triple Negative, 1 ER^+^ and 1 HER2^+^) were implanted in the brain parenchyma following the same procedure as for the 436 cells. The tumor intake rate was greater than 88% for each sample, establishing the developing brain parenchyma as an environment of choice for breast tumor cell survival and growth (**Figures 6A and 6D**). Furthermore, depending on the patient sample, we noted different tumorigenic behaviors, ranging from dense, localized tumor masses to numerous small tumor foci implanted in the parenchyma or floating in the cerebrospinal fluid (**Figure 7D**). Thus, we succeeded with our *in vivo* model to establish various metastatic models that allow tumor cells to be subjected to different microenvironments and therapeutic regimens.

**Figure 6:**
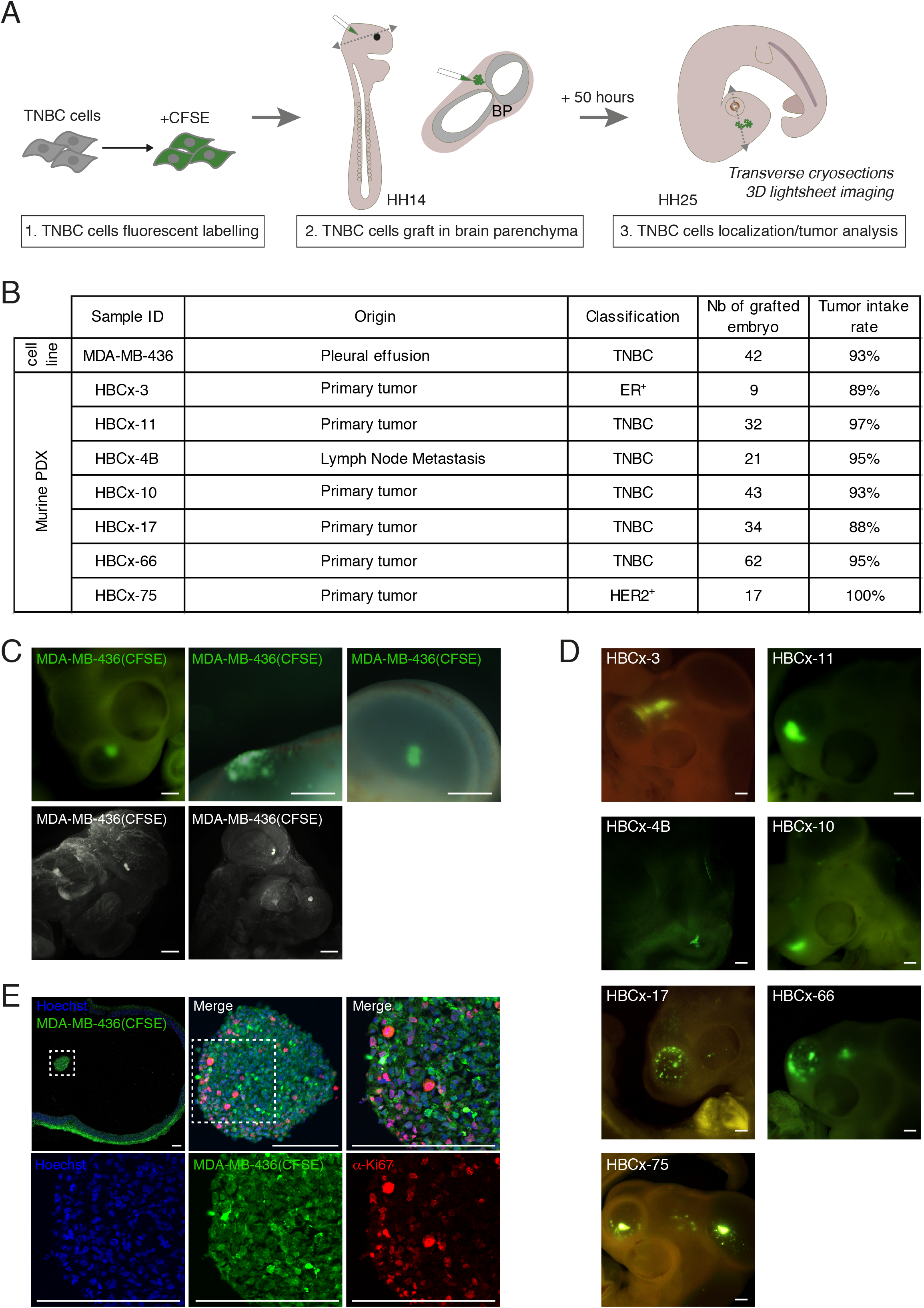
Targeting the developing brain to model TNBC cerebral metastasis in the avian embryo. **A**. Set up of TNBC cells grafting procedure in the developing brain parenchyma (BP): TNBC cells are labeled with CFSE and microinjected in the developing BP at HH14 stage. Grafted embryos are harvested for imaging analysis 48 hours after the graft, at HH25 stage. **B**. Table recapitulating breast cancer samples characteristics – origin, classification – and their corresponding tumor intake rate in the avian BP. **C**. Representative images of classical localization of MDA-MB-436 cells in the developing head 48 hours after the graft as observed with the stereomicroscope (upper panels) or by 3D lightsheet imaging (lower panels). Scale bar: 300 µm. **D**. Detection of Ki67+ proliferating cells by immunofluorescence (in red) on cryosections of MDA-MB-436 tumor masses (in green, CFSE+) formed in the developing brain of chick embryos. Chick and human nuclei are labelled with Hoechst (in blue). Scale bars: 100 µm. **E**. Representative images of breast cancer patient cells in the developing head 48 hours after the graft as observed with the stereomicroscope. Scale bars: 300 µm.

## Discussion

Our study reports a novel *in vivo* paradigm to model breast cancer tumorigenesis and applications of this model to various preclinical investigations. When implanted in targeted tissues of the avian embryo, human TNBC cells from diverse origins such as cell lines, murine PDX, patient biopsies and resections (from both primary and metastatic foci) were all fully capable of surviving, proliferating and forming tumors in a rapid and reproducible manner. As we expected, the somitic region committed to becoming skeletal tissues provides a favorable microenvironment to TNBC cells. Notably, none of 11 engrafted patient samples resulted in failure of tumor intake, with a greater than 90% success rate for 8 of them. This makes our innovative model ideally suited for statistical analysis of tumor volumes and quantitative comparison of experimental conditions. Administration of therapeutic compounds to grafted embryos and subsequent 3D analysis of tumor volumes revealed that with our process, drug effects can be detected in as little as one day post-treatment, which opens new avenues for rapid screening therapeutic strategies. We also demonstrated that our *in vivo* model is adapted to the sorting of miniature tumors, and the running of large-scale molecular analysis on sorted tumor cells, which thus enable deep characterization of the mechanisms of action of therapeutic compounds.

Our results with gemcitabine and carboplatin (standards of care for TNBC patients) are consistent with those reported with murine models and in the clinic (Karginova et al., 2015; Maisano et al., 2011; Ou et al., 2020).

First, we could reproduce the responsiveness of MDA-MB-436 cells to gemcitabine and carboplatin (Karginova et al., 2015; Sasaki et al., 2014).

Second, our transcriptomic analysis comparing tumors formed in the avian tissues that were exposed to the gemcitabine/carboplatin combination and excipient revealed a landscape of gene pathway regulation that is fully consistent with the mode of action of these chemotherapies. Similar to other platinum derivatives, carboplatin covalently binds to the N7 site of purine bases, interfering with cell replication, which drives the cells towards apoptosis or necrosis (Schoch et al., 2020). Gemcitabine is an analog of deoxycytidine and its active, phosphorylated form interferes with DNA synthesis (Plunkett et al., 1995). Of the top gene pathways identified, most were related to DNA damage response and cell death. These findings demonstrate that 24 hours’ exposure of engrafted tumor cells to these chemotherapies through delivery via the general circulation reproduces their expected outcome. This demonstrates the strength of both our *in vivo* model and its companion procedure for RNAseq analysis of chemotherapy-treated tumors grown in the avian embryo.

Third and interestingly, when the avian model was applied to 4 patient samples, we observed for half of them that their avian tumor replicas were significantly reduced by SOC administration, with one case showing a particularly pronounced response. For the remaining half, no significant effect of chemotherapy administration was observed. These findings reflect acknowledged heterogeneity among TNBC patient tumors, with a gene profiling-based stratification of TNBCs into several sub-types, which is translated into different responses to chemotherapies (Gupta et al., 2020; Lehmann et al., 2011). Heterogeneity was also reflected in our paradigm of implantation in the avian embryo brain parenchyma. We observed various behaviors of tumor cells depending on patients, with variable propensity of cells to widely colonize the brain tissues. In some cases, tumor cells adopted a constellation pattern whereas in others they could form either a dense mass or display a mixed pattern. Interestingly too, ER+ and HER2+ tumor cells were as capable as more aggressive triple negative ones of forming tumors. This indicates that the avian brain microenvironment provides effective support for tumorigenesis while also allowing heterogeneity between patients to be manifested.

Our data with ASNase supports the relevance of using of our approach for preclinical investigation. Anti-tumor efficacy was demonstrated for ASNase in both the MDA-MB-436 TNBC cell line and patient biopsies, and suited for evaluating both therapeutic compounds at different doses and combination therapies. This key preclinical finding strongly substantiates the case for advancing ASNase therapies to TNBC clinical trials, and provided a rational for the ongoing study in metastatic TNBC, in which eryaspase (ASNase encapsulated in red blood cells) is evaluated in combination with gemcitabine and carboplatin (NCT03674242).

Altogether, these findings establish our *in vivo* model of tumoral cell implantation in targeted regions of the avian embryo as a model of choice for preclinical investigations, conveniently integrating patient stratification in the therapeutic efficacy evaluation process, which is a major asset for the advancement to clinical trials. Furthermore, coupling the generation of patient tumor replicas to large-scale molecular analysis will open new avenues for the characterization of mechanisms of action of therapeutic compounds, the prediction of their optimal combinations and the development of personalized medicine.

## Acknowledgments

We thank Alexander Scheer, Françoise Horand, Marie Châlons-Cottavoz and Bastien Laperrousaz for helpful discussions, Priscillia Rebus and Florian Tavernier for *in vitro* experiments performed at Erytech Pharma, Oncodesign SAS for the sharing of TNBC samples from their PDX murine models and associated clinical information, and ProfileXpert for providing advice on RNAseq data analysis. We thank Gil Beyen, Iman El-Hariri, Dan Cole, Chad Kitchen, Nigel Biswas-Baldwin (from Erytech) and Sabine Poppenborg (from medac GmbH) for the review of this article and their useful advice.

## Material and Methods

### Cell lines

The MDA-MB-436 TNBC cell line was purchased by Erytech Pharma and licensed from the M.D. Anderson Cancer Center (agreement ID: OCT20-002). The cell line was cultivated in Roswell Park Memorial Institute Medium (RPMI 1640, Gibco) supplemented with 10% Fetal Bovine Serum (FBS) and 25 U/mL Penicillin Streptomycin (Sigma).

### Patient samples

Patient samples OF-BRE-nbt219, OF-BRE-nbt1111, OF-BRE-nbt311, OF-BRE-nbt783, OF-BRE-nbt494 and associated data were obtained from NeuroBioTec (CRB HCL, Lyon France, Biobank BB-0033-00046) and are part of a collection declared at the French Department of Research (DC 2008-72).

Patient samples OF-BRE-001, OF-BRE-002, OF-BRE-007, OF-BRE-012, OF-BRE-016, OF-BRE-017 were obtained from Oncofactory’s collection declared at the French Department of Research (DC-2018-3275). Patient informed consent was obtained according to French ethical rules for each sample, and collected by referent hospital practitioners (Hopital Jean Mermoz, Lyon France; Hospices Civils de Lyon, France).

Patient-Derived Xenografts (PDX) OD-BRE-0503, OD-BRE-0589, OD-BRE-0631, OD-BRE-0733 were provided by Oncodesign and established from triple negative breast cancer patient samples by Imodi platform (www.imodi-cancer.com).

Patient-Derived Xenografts (PDX) HBCx-3, HBCx-11, HBCx-4B, HBCx-10, HBCx-17, HBCx-66, HBCx-75 were established from breast cancer patients with informed consent from the patient in accordance with published protocols (Marangoni et al. CCR 2007; Marangoni et al. CCR 2018 ; Coussy et al. IJC 2019).

### Anticancer drugs

Gemcitabine (stock solution: 100 mg/mL) and carboplatin (stock solution: 10 mg/mL) were purchased from Accord Healthcare and diluted in NaCl 0.9% for *in vivo* experiments.

L-Asparaginase (ASNase) was purchased from medac GmbH (Spectrila®, 81021-K19, batch: D150857C) and resuspended in sterile NaCl 0.9% for *in vivo* experiments.

### *In vitro* cell survival assay

For *in vitro* cytotoxicity assays, 2500 MDA-MB-436 cells were plated in 96-well plates. Treatment with increasing doses of ASNase was performed 24 hours after plating. Cell survival measurement was performed after 4 days of treatment, using a cell counting Kit-8 colorimetric assay (CCK-8 kit, Sigma, 96992). Each experiment was repeated three times. IC50 calculation was performed using Prism 8.0 (GraphPad) software.

### *In ovo* xenograft

Embryonated eggs were obtained from a local supplier (Couvoir de Cerveloup, Vourey, France) and incubated at 38.5°C in a humidified incubator. Cell lines or patient samples were dissociated and labeled with 8 µM CFSE solution (Life Technologies). Stage HH14 chick embryos were grafted with fluorescent cells in presumptive somitic areas or in the brain parenchyma, with a glass capillary connected to a pneumatic PicoPump (PV820, World Precision Instruments) under a fluorescence stereomicroscope.

### Evaluation of drug toxicity in chick embryos

Twenty-four (24) hours after drug intravenous injection, chick embryos were harvested, weighted (Sartorius Quintix35-1S) and measured along the rostro-caudal axis using Leica LASX image analysis software. The Body Surface Area (BSA) was calculated using Dubois & Dubois formula: BSA (m^2^)= 0.20247 x height (m)^0.725^ x weight (kg)^0.425^.

The morphology / anatomy of each embryo was systematically analyzed to check their correct stage-related development. The criteria observed were: the survival (heart beating), the craniofacial morphology (presence of each cerebral compartment and eyes), the presence of four limb buds, the cardiac morphology, and the anatomy of embryonic annexes such as the allantois.

### Immunofluorescence on cryosections

Chick embryos were harvested and fixed in 4% Paraformaldehyde (PFA). Embryos were embedded in 7,5% gelatin-15% sucrose in PBS to perform 20 µm transverse cryosections. Permeabilisation and saturation of sections was performed in PBS-BSA 3%-Triton 0.5%. Anti-Ki67 (1/200, ab15580, Abcam) primary antibody was applied to cryosections. Alexa 555 anti-rabbit IgG (1/500, A21206, Life Technologies) was used as secondary antibody. Nuclei were stained with Hoechst (H21486, Invitrogen). Slices were imaged with a confocal microscope (Olympus, FV1000, X81) using either a 10X objective for whole slice imaging or a 40X objective to focus on Ki67 immunolabeling.

### Tissue clearing and whole mount SPIM imaging

PFA-fixed HH25 embryos were cleared using an adapted Ethyl-Cinnamate protocol (Klingberg et al, 2017). Briefly, tissues were dehydrated in ethanol successive baths finally cleared in Ethyl Cinnamate (Sigma, 112372). Cleared samples were imaged using the UltraMicroscope SPIM (LaVision Biotech). 3D-images were built using Imaris™ software. Volumetric analysis was performed using Imaris™ “ Surface” module adjusted on CFSE fluorescence.

### Statistical treatment of the data

Statistical treatment of the data was performed with Prism 6.0e (GraphPad). For parametric tests, both normality and variances homoscedasticity were checked. All statistical tests were two-sided.

### Concentration of plasma amino-acids by HPLC

Collected blood samples underwent a centrifugation process (1000g, 4°C for 10min) to prepare plasmas. Collected supernatants (plasmas) were aliquoted and stored at -20°C until the dosage of amino acids. The determination of amino acids level (ASN, GLN, ASP, GLU) in chicken embryo plasma was performed by Ion-pairing reversed-phase liquid chromatography coupled to tandem mass spectrophotometry (HPLC/MSMS) method. Chicken embryo plasma was considered depleted in ASN when concentration is < 2 μM demonstrating the efficacy of ASNase activity (depending on doses tested).

### RNA isolation and library preparation for high-throughput mRNA sequencing (RNA-seq)

MDA-MB-436 tumors developed in chick embryos treated either gemcitabine/carboplatin or excipient were microdissected 48 hours after grafting using a stereomicroscope. Dissected tissues were dissociated using an enzymatic cocktail comprising 1.25 mg/mL Collagenase IV, 50 µg/mL DNAse and 2.9 mg/mL trypsin-EDTA. CFSE+ cells were sorted using CellenONE cell sorter (Cellenion). The experiment was performed in duplicates for each condition. The experimental design thus consisted of 4 samples divided into 2 conditions: Excipient and gemcitabine/carboplatin. RNA isolation from sorted cells and RNA-Seq processing was performed at the ProfileXpert core facility (Lyon, France). Total RNA was extracted with Norgen single cell RNA purification kit (Norgen) and the quality was checked with a Bioanalyzer 2100 (Agilent, RIN >8.0). Ribosomic RNA was depleted with Ribo-Zero™ Gold Kit (Epicentre). RNA-seq libraries were prepared with SMART-Seq® Stranded Kit (Takara-Clontech). Samples were sequenced in wholeRNAseq using the NextSeq illumina 500 Platform (75 bp single read). Reads were mapped using TopHat v.2.1.0 against the human Genome build (hg38). Quantification was done using HTSeq-count software (0.11.3). The differential analysis was performed with the DESeq2 tool (3.11) with median of ratios normalisation that allows comparison between samples.

A gene is considered to be upregulated or downregulated when its level of expression varies with a fold change greater than or equal to 1.5 between the two conditions and its adjusted p-value (padj) is less than 0.05. Transcripts with detectable expression in one condition (ON) versus no detectable expression in the other condition (OFF) were also added to the list of differentially expressed transcripts.

Analysis of main functions concerned by gene expression change was performed with ToppFun (https://toppgene.cchmc.org/enrichment.jsp)

## Notes

### Competing Interest Statement

The authors have declared no competing interest.

